# Transition Metal Dichalcogenide Nanoflowers Rescue Immune Cells from the Cytotoxic Effects of Amyloid Aggregates

**DOI:** 10.1101/2025.10.02.680126

**Authors:** Mikhail Matveyenka, Charles L. Mitchell, Dmitry Kurouski

**Author notes:** **Corresponding Author** *Dmitry Kurouski*.

## Abstract

Parkinson’s disease (PD) is a severe pathology caused by a progressive degeneration of neurons in the substantia nigra pars compacta, hypothalamus, and thalamus. Although etiology of PD remains unclear, accumulating evidence indicates that neurodegenerative effects are triggered by the abrupt aggregation of α-synuclein (α-Syn), a small membrane protein that is responsible for cell vesicle trafficking. α-Syn aggregates are highly toxic to neurons and immune cells present in the brain, including macrophages, microglia, and dendritic cells. Transition metal dichalcogenide nanoflowers (TMD NFs) are novel nanomaterials with unique optical and biological properties. However, their effects on the immune system remain poorly understood. In this study, we investigate cytoprotective properties of molybdenum disulfide (MoS_2_) and molybdenum diselenide (MoSe_2_) NFs on macrophages, microglia, and dendritic cells exposed to α-Syn fibrils. We found that MoSe_2_ NFs exerted strong cytoprotective properties fully mitigating toxic effects of α-Syn fibrils, while MoS_2_ NFs were found to be significantly less potent in rescuing immune cells from α-Syn aggregates. At the same time, MoS_2_ NFs triggered polarization of macrophages into M1 and dendritic cells into M2 phenotypes, while an increase in both M1 and M2 was observed in microglia exposed to MoS_2_ NFs. MoSe_2_ NFs did not trigger polarization of DC cells and microglia in M1/M2 phenotypes, while MoSe_2_ NFs-facilitated polarization of macrophages into M1 was observed. These results indicate that TMD NFs could be used to improve viability of immune cells and attenuate their phenotypes, which, ultimately, can be used to treat PD and other neurodegenerative pathologies.

## Introduction

Parkinson’s disease (PD) is a rapidly growing neurodegenerative pathology that is expected to strike 12 million patients by 2040 worldwide.^1^ In the US alone, over 60,000 new PD diagnoses are reported annually, putting an enormous burden on our society.^2^ All currently available therapeutics only alleviate PD symptoms, making a neuroprotective treatment an urgent and unmet need.^3^

On the molecular level, PD is characterized by the progressive aggregation of α-synuclein (α-Syn), a small protein localized in the cell membranes, which results in the formation of oligomers and fibrils.^4–12^ These protein aggregates exert cytotoxic effects to neurons primarily localized in the midbrain, hypothalamus, and thalamus, as well as other brain areas.^13–23^ Experimental findings reported by our group show that α-Syn oligomers and fibrils are endocytosed by neurons which results in irreversible damage of cell endosomes.^24, 25^ As a result, α-Syn aggregates leak into the cytosol where they impair mitochondria and endoplasmic reticulum, triggering the unfolded protein response. These and other cytotoxic effects trigger an increase in the levels of reactive oxygen species (ROS) and ultimately cause neuronal death.

Accumulating evidence indicates that the onset and progression of PD can be linked to α-Syn-triggered degeneration of brain immune cells, including macrophages, dendritic cells, and microglia.^26–29^ Macrophages and microglia endocytose and digest viruses and microorganisms, as well as degrade apoptotic cells.^30, 31^ Dendritic cells have similar activities. However, they retain the endocytosed antigens and present them on the major histocompatibility complex (MHC)^32^, as part of the adaptive immune system. Antigen presentation by dendritic cells activates T cells, as well as triggers other cascades required for the activation of the immune system to maintain the homeostasis of the central nervous system.^26–29^ Thus, α-Syn-triggered degeneration of immune cells weakens brain defense.^27–29^ It was shown that amyloid aggregates facilitate polarization of macrophages into pro-inflammatory (M1) phenotype, which triggers brain inflammation that accelerates neurodegeneration.^33^

Recently our group demonstrated that transition metal dichalcogenide nanoflowers (TMD NFs) exerted neuroprotective effects in neurons and astrocytes.^34^ These neuroprotective effects are triggered by NFs-induced mitochondrial biogenesis and upregulation of peroxisome proliferator-activated receptor gamma coactivator 1-alpha (PGC-1α).^34^ However, it was unclear whether TMD NFs exert any cytoprotective effect to immune cells exposed to α-Syn aggregates. In the current study, we employed a set of molecular and cell assays to investigate the extent to which molybdenum disulfide (MoS_2_) and molybdenum diselenide (MoSe_2_) NFs alleviate cytotoxic effects of α-Syn fibrils. We also examined dynamics of TMD NFs-induced polarization of macrophages, dendritic cells, and microglia into pro- (M1) and anti-inflammatory (M2) phenotypes.

## Results

We utilized cell proliferation assay to determine whether TMD NFs alone have any cytotoxic effects on the immune cells. We found that at low concentrations (1.5-25µg), both MoSe_2_ and MoS_2_ NFs facilitated cell proliferation. At higher concentrations (50µg), a small decrease in the proliferation of macrophages was observed in the presence of MoS_2_ NFs, whereas no effect was evident for macrophages exposed to MoSe_2_ NFs, Figure S1. DC cells and microglia exhibited a gradient decrease in proliferation when exposed to increasing concentrations of MoS_2_ NFs, while this negative effect was not observed in the cells treated with MoSe_2_ NFs, Figure S1. These results indicate that a concentration range of 25-50µg is optimal for MoSe_2_ NFs, whereas MoS_2_ NFs are slightly less biocompatible at these concentrations.

Using ROS assay, we determined the extent to which macrophages, dendritic cells, and microglia exposed to α-Syn fibrils could be rescued by TMD NFs. We found that α-Syn fibrils triggered an increase in the ROS levels in macrophages, DC, microglia, and neurons, Figure 1. At the same time, administration of MoSe_2_ NFs at 50µg at 12h after cell exposition to α-Syn fibrils fully rescued all cell types from the cytotoxic effect of amyloid fibrils. It should be noted that MoS_2_ NFs exerted cytoprotective properties only in DC cells and neurons, while no effects of MoS_2_ NFs activity were observed in macrophages and microglia. These results indicate that MoSe_2_ NFs fully rescued different immune cells from the cytotoxic effects of α-Syn aggregates, while MoS_2_ NFs were found cytoprotective only for DC cells and neurons. Thus, MoSe_2_ and MoS_2_ NFs have dissimilar cytoprotective properties in different cell types.^34^

**Figure 1.**
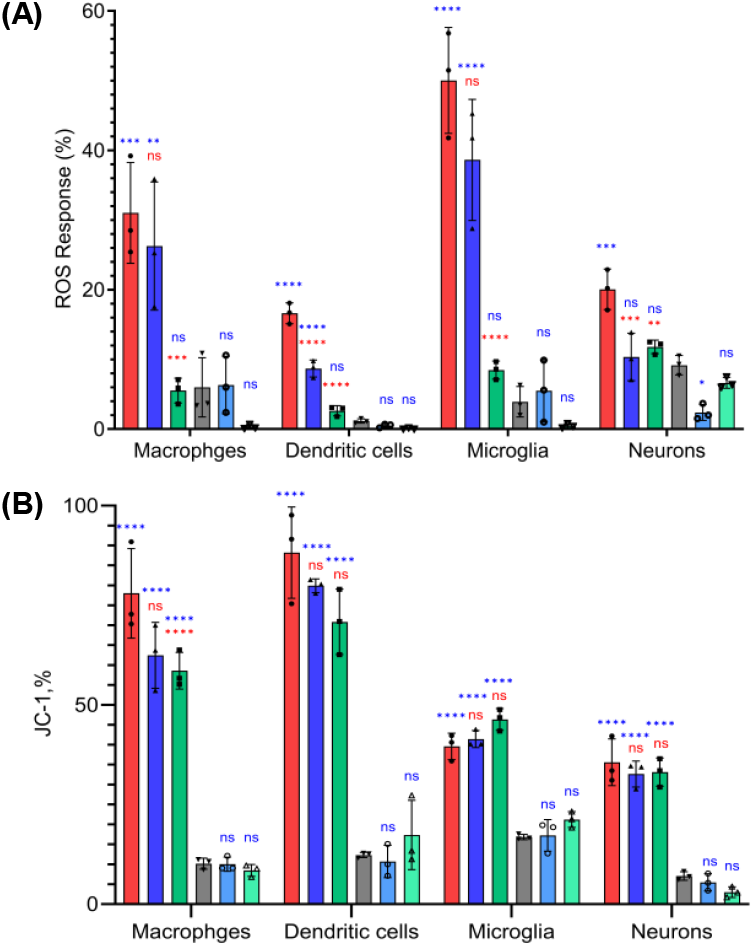
Bar graphs of ROS (A) and JC-1 (B) assays showing response of macrophages, dendritic cells, microglia, and neurons to α-Syn fibrils (red), α-Syn fibrils with subsequent administration of 50 µM of MoS_2_ NFs (blue), and MoSe_2_ NFs (green), as well as MoS_2_ NFs (light blue) and MoSe_2_ NFs (light green) alone. Control cells with PBS added are shown in grey. According to ANOVA, * P< 0.05, ** P< 0.005, *** P< 0.0005, **** P< 0.0001 and NS – not significant. Blue significance markings relate to comparisons made to control cells that received neither α-syn fibrils nor TMD NFs. Red significance markings relate to comparisons made to cells that were exposed to α-syn fibrils.

It should be noted that in macrophages, microglia, and DC not exposed to α-Syn fibrils, administration of MoSe_2_ NFs fully removed cytosolic ROS, while a strong suppression of ROS levels was observed in neurons exposed to these nanomaterials. The magnitude of such ROS suppression activity was significantly lower for MoSe_2_ compared to MoS_2_ NFs. It is important to note that ROS levels in all cells exposed to MoS_2_ NFs were similar to the control, which indicates that poor cytoprotective properties of MoS_2_ NFs discussed above unlikely can be caused by their cytotoxic effects at this concentration (50 µg).

One can expect that ROS suppression activity of MoSe_2_ NFs is provided by superoxide dismutase properties of the nanomaterials.^35–37^ In this case, Mo atoms located in the corners, edges, and surface defects of MoSe_2_ NFs can change redox states from Mo^+6^ to Mo^+4^ and vice versa, which will result in the degradation of ROS in the cells (i and ii).

i. Mo^4+^ + O_2_^•-^ + H^+^ → Mo^6+^ + H_2_O_2_,
ii. Mo^6+^ + O_2_ ^•-^ → Mo^4+^ + O_2_ + H^+^.

We also anticipate that MoSe_2_ NFs exert catalase-like activity which allows for the removal of hydrogen peroxide in the cell cytosol (iii and iv).

(iii) Mo^6+^ + H_2_O_2_ + 2OH^-^ → Mo^4+^ + O_2_ + 2H_2_O,
(iv) Mo^4+^ + H_2_O_2_ → Mo^6+^ + 2OH^-^.

Previously, our group demonstrated that endocytosis of amyloid fibrils triggers damage of cell endosomes.^38–40^ As a result of such damage, amyloid fibrils leak into the cytosol where they impair mitochondria and other organelles.^24, 41^ Expanding upon this, we investigate whether TMD NFs can alleviate amyloid-induced impairment of mitochondria in the immune cells. Using JC-1 assay, we observed a significant decrease in the degree of mitochondrial impairment in macrophages treated with α-Syn fibrils and MoSe_2_ NFs compared to the cells exposed to α-Syn fibrils alone. However, cell treatment with MoS_2_ NFs resulted only in the insignificant decrease in the magnitude of mitochondrial impairment triggered by α-Syn fibrils. Similar observations were made for other cell types. These results indicate that TMD NFs are likely to both directly protect mitochondria in the immune cells and enact cytoprotective properties via different cell mechanisms, including degradation of ROS and hydrogen peroxide as discussed above.

Next, we investigated whether TMD NFs could polarize microglia towards M1 and M2 phenotypes. For this, changes in the expression of CD86 and CD206 receptors were analyzed, Figure 2 and Table 1. We found that α-Syn fibrils triggered polarization of macrophages towards M1 phenotype. Administration of MoSe_2_ and MoS_2_ NFs afterwards did not significantly change the ratio between M1 and M2 populations. Interestingly, administration of TMD NFs to the cells with no α-Syn fibril exposure also did not increase the amount of M1 population compared to the control. However, a substantial decrease in M2 population was observed in such cases. Thus, we can conclude that TMD NFs alter the ratio between macrophage M1 and M2 populations but did not increase the percentage of M1 cells.

**Table 1.**
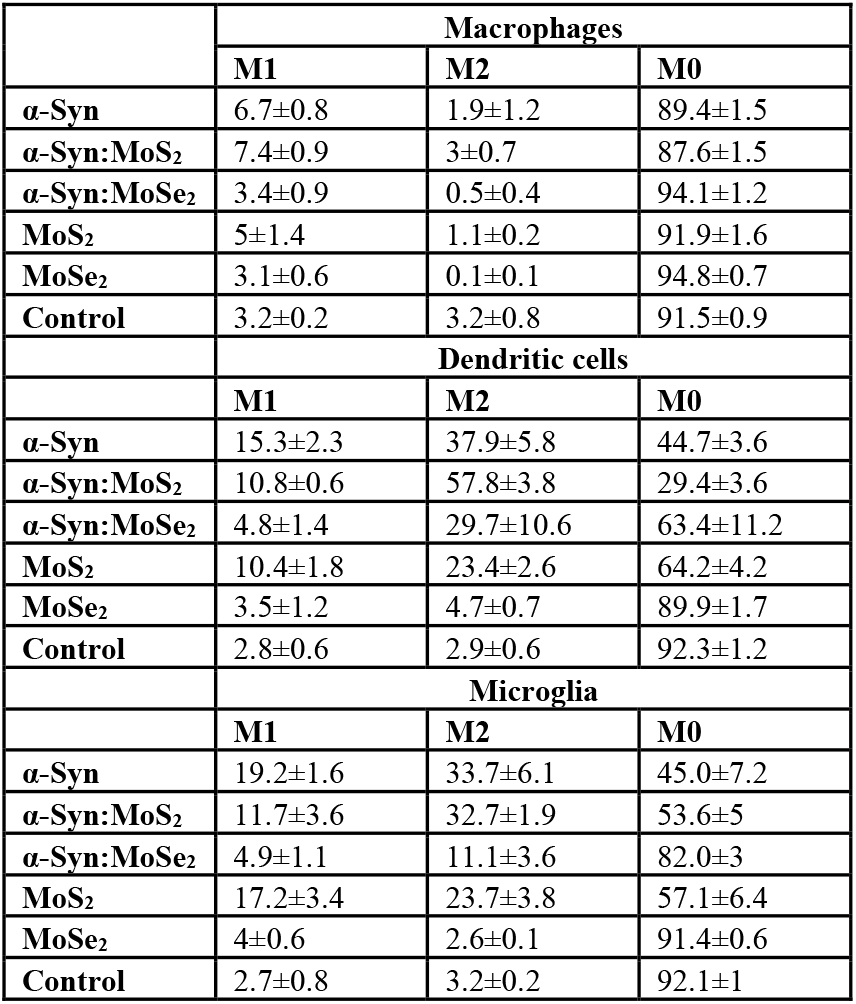
M1, M2, and M0 population percentages of macrophages, dendritic cells, and microglia after administration of α-Syn fibrils (α-Syn) alone and with the subsequent (12h later) treatment with 50 µM of MoS_2_ (α-Syn:MoS_2_) and MoSe_2_ NFs (α-Syn:MoSe_2_), as well as cells treated with MoS_2_ (MoS_2_) and MoSe_2_ NFs (MoSe_2_) alone (50 µM).

**Figure 2.**
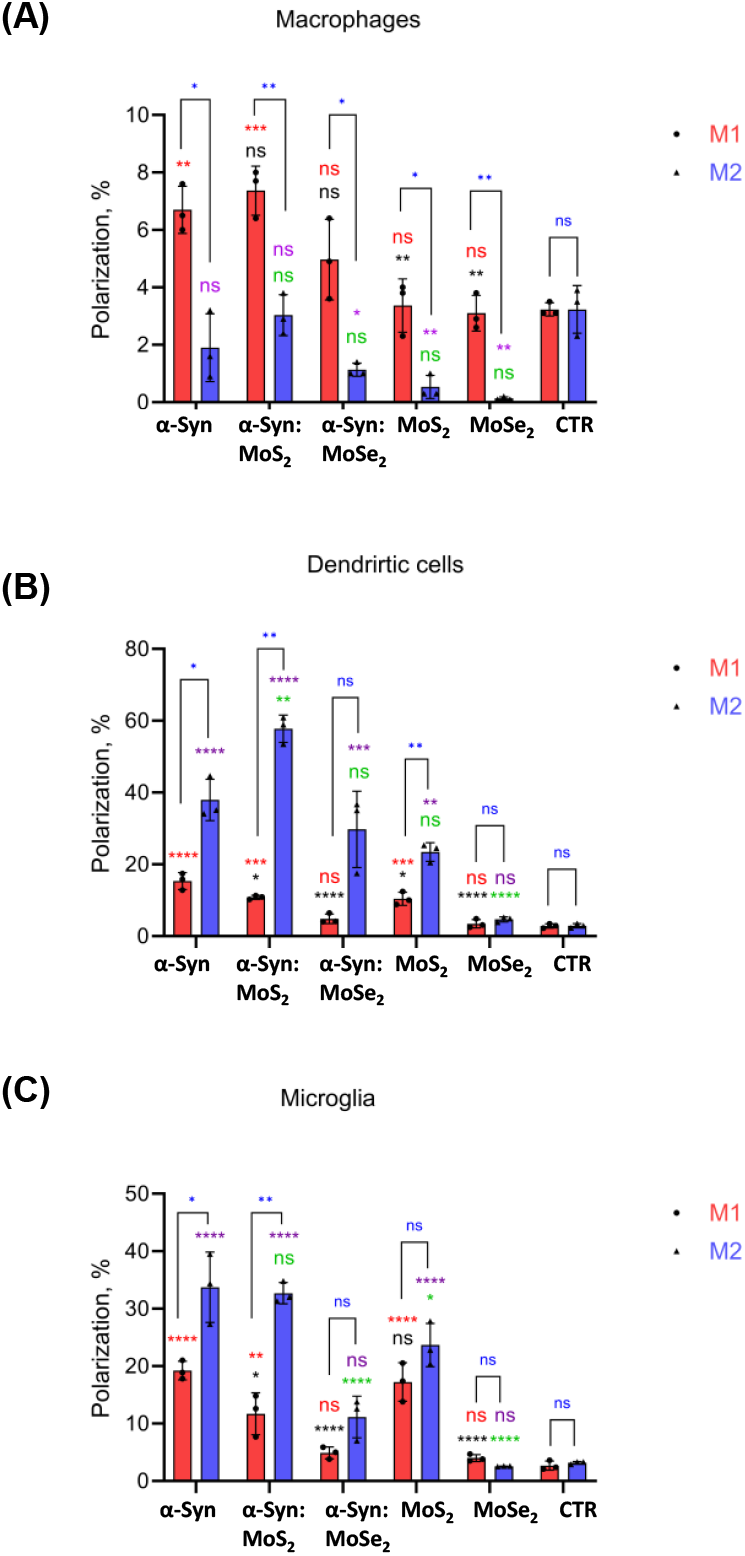
Bar graphs of M1 (CD86) and M2 (CD206) populations of macrophages, dendritic cells, and microglia after administration of α-Syn fibrils (α-Syn) alone, and with the subsequent (12h later) treatment with 50 µM of MoS_2_ (α-Syn:MoS_2_) and MoSe_2_ NFs (α-Syn:MoSe_2_), as well as cells treated with MoS_2_ (MoS_2_) and MoSe_2_ NFs (MoSe_2_) alone (50 µM). According to ANOVA, * P< 0.05, ** P< 0.005, *** P< 0.0005, **** P< 0.0001 and NS – not significant. Red (M1) and purple (M2) significance markings relate to comparisons made to control cells that received neither α-syn fibrils nor TMD NFs. Black (M1) and green (M2) significance markings relate to comparisons made to cells that were exposed to α-syn fibrils. Blue significance markings relate to comparisons made between M1 and M2.

We also found that MoS_2_ NFs triggered polarization of DC cells towards M2 phenotype, while this effect was not observed for MoSe_2_ NFs, Figure 2 and Table 1. α-Syn fibrils also triggered polarization of DC cells into M2. Subsequent administration of MoS_2_ NFs to such cells further increased the amount of M2 population and decreased M1 population. Similar effect was observed for MoSe_2_ NFs. These nanostructures drastically lowered the amount of M1 DC cells, while no significant changes in the amount of M2 cells were observed compared to DC cells exposed to α-Syn fibrils.

Our results show that α-Syn fibrils polarize microglia towards M2 phenotype, Figure 2 and Table 1. MoS2 NFs slightly lowered the amount of M1 population if subsequently administered to the cells, whereas no effect on M2 population was observed. Administration of MoSe_2_ NFs, on the contrary, drastically lowered the amount of both M1 and M2 microglia. It should be noted that MoSe_2_ NFs alone did not change microglia polarization, while a substantial increase in both M1 and M2 was observed for MoS_2_ NFs.

## Conclusions

Our results demonstrate that MoSe_2_ NFs can rescue immune cells from cytotoxic effects of α-Syn fibrils. MoS_2_ NFs exerted substantially weaker cytoprotective properties under the same experimental conditions. We also demonstrated that TMD NFs facilitate proliferation of the immune cells into M1 and M2 phenotypes. These results suggest that MoSe_2_ NFs can be utilized to suppress microglia-induced brain inflammation, as well as facilitate wound healing, and treat muscle and spinal cord injuries.

## Experimental Section

### Protein expression and purification α-syn

Rosetta strain of *Escherichia coli* BL21 (DE3) was transformed with pET21a-α-synuclein plasmid. Next, bacterial cells grew in the media to reach an optical density of 0.8. After that, 1 mM of isopropyl-ß-D-thiogalactopyranoside (IPTG) was added to the media to trigger the protein expression for 4-5 h. Cells were harvested by centrifugation at 8, 000 rpm for 10 min. The pellet was resuspended in lysis-tris buffer (50 mM Tris, 10 mM EDT, 150 mM NaCl, pH 7.5). The suspended cells placed in boiling water 30 min to complete cell lysis and centrifuged at 16,000 g for 40 min. The supernatant enriched with α-Syn was collected. Streptomycin sulfate (10% solution, 136 μL/mL) in glacial acetic acid (228 μL/mL) was added to the supernatant to precipitate bacterial proteins and nucleic acids. Final solution was then centrifuged at 16,000 g for 10 min at 4 °C. Supernatant was precipitated by saturated ammonium sulfate ((NH_4_)_2_SO_4_) at 4 °C. Samples were then centrifuged to collect α-Syn pellet that was subsequently washed one more time with (NH_4_)_2_SO_4_ at 4 °C (a 1:1 v/v mixture of saturated (NH_4_)_2_SO_4_ and water). Next, the pellet was re-suspended in 100 mM NH_4_(CH_3_COO) under constant stirring for 5 min and absolute ethanol was added to the resuspended α-Syn to precipitate the protein. This step was repeated twice at room temperature. Finally, collected protein were dissolved in sterile PBS.

### Size Exclusion Chromatography (SEC)

Protein solutions were first centrifuged at 14,000 g for 30 min using a benchtop microcentrifuge (Eppendorf centrifuge 5424 USA). Next, 500 μL of the concentrated protein was injected into an AKTA pure (GE Healthcare) FPLC system equipped with a gel filtration column (Superdex 200 10/300). Samples were eluted isocratically using PBS, pH 7.4 at a flow rate of 0.5 mL/min. All protein purification was done at 4 °C. During the FPLC run, (iii) mL fractions were collected based on UV-VIS detection at 280 nm. Collected fractions of α-Syn were concentrated via 10 kDa Amicon Ultra protein filters (Merck Millipore).

### Protein aggregation

α-Syn was diluted in PBS to reach the final protein final concentration of 400 μM and incubated in Eppendorf tubes at 37 °C for 120 h under constant 510 rpm agitation.

### Cell culturing and CD86/CD206 assay

Murine macrophage-like (RAW264.7), Dendritic (DC2.4), and Microglia (SIM-A9) cell lines were obtained from American Type Culture Collection (ATCC); mycoplasma testing was negative. Cells were maintaned under standard conditions in RPMI-1640 (Corning, USA) with 10% of FBS and antibiotic (Gibco, Waltham, MA, USA); SIM-A9 microglial cells were maintained in DMEM/F-12 (Gibco, Waltham, MA, USA) supplemented with 8% FBS, 4% horse serum (HS) (Corning, USA) and antibiotic. All cell lines were seeded in 96-well plates at a predefined density and allowed to attach at 37°C, 5% CO_2_. On the following day, α-Syn (40 µM) was added and cultures were incubated for 12h. After 12h, nanoflowers (NFs) were added and cells were incubated for an additional 24h. Cells were collected, washed twice with staining buffer (PBS, 2% FBS). Then labeled and incubated with fluorophore-conjugated anti-CD86 (cat. #105021, BioLegend) and anti-CD206 (cat. #141707, BioLegend) at final concentration of 0.5 μg per 10^6^ cells in 100 µL (i.e., 5 µg/mL) for 30 min at 4°C in the dark. The cells were washed and resuspended in 100 µL PBS. Data was acquired on a multicolor flow cytometer BD LSR II equipped with blue, red, and violet lasers. Analysis proceeded through debris exclusion live-cell gating, followed by lineage identification where applicable. CD86 versus CD206 bivariant plots were used to quantify M1-like (CD86^high^ / CD206^low^), M2-like (CD86^low^ / CD206^high^), and intermediate populations.

### ROS assay

For the Reactive oxygen species (ROS) Assay, cell lines were prepared as described above. Cells were washed with PBS, stained with CellROX reagent (Invitrogen, Waltham, MA, USA) (final concentrtion 5 µM) for 30 min at 37°C, washed with PBS, and analyzed on a BD LSR II flow cytometer.

### JC-1 assay

For the JC-1 Assay (Invitrogen, Waltham, MA, USA), cells were prepared as described above, washed with PBS, incubated with JC-1 (2 µg/mL) for 30 min 37°C, protected from light, rewashed, and acquired on a BD LSR II. Green monomers were collected on FITC channel (530/30), and red aggregates on PE channel (585/42), and data were expressed as the red/green fluorescence ratio.

## Notes

The authors declare no competing financial interests

## ACKNOWLEDGMENT

We are grateful to the National Institute of Health for the provided financial support (R35GM142869).

## REFERENCES

(1) Chen, J. Parkinson’s disease: health-related quality of life, economic cost, and implications of early treatment. J. Am. J. Manag. Care 2010, 16, S87–S93.

(2) Bengoa-Vergniory, N.; Roberts, R. F.; Wade-Martins, R.; Alegre-Abarrategui, J. Alpha-synuclein oligomers: a new hope. Acta Neuropathol 2017, 134 (6), 819–838. DOI: 10.1007/s00401-017-1755-1.

(3) Shahmoradian, S. H.; Lewis, A. J.; Genoud, C.; Hench, J.; Moors, T. E.; Navarro, P. P.; Castano-Diez, D.; Schweighauser, G.; Graff-Meyer, A.; Goldie, K. N.; et al. Lewy pathology in Parkinson’s disease consists of crowded organelles and lipid membranes. Nat Neurosci 2019, 22 (7), 1099–1109. DOI: 10.1038/s41593-019-0423-2.

(4) Spillantini, M. G.; Schmidt, M. L.; Lee, V. M.; Trojanowski, J. Q.; Jakes, R.; Goedert, M. Alpha-synuclein in Lewy bodies. Nature 1997, 388 (6645), 839–840. DOI: 10.1038/42166.

(5) Baba, M.; Nakajo, S.; Tu, P. H.; Tomita, T.; Nakaya, K.; Lee, V. M.; Trojanowski, J. Q.; Iwatsubo, T. Aggregation of alphasynuclein in Lewy bodies of sporadic Parkinson’s disease and dementia with Lewy bodies. Am. J. Pathol. 1998, 152 (4), 879–884.

(6) Davidson, W. S.; Jonas, A.; Clayton, D. F.; George, J. M. Stabilization of alpha-synuclein secondary structure upon binding to synthetic membranes. J Biol Chem 1998, 273 (16), 9443–9449. DOI: 10.1074/jbc.273.16.9443.

(7) Kruger, R.; Kuhn, W.; Muller, T.; Woitalla, D.; Graeber, M.; Kosel, S.; Przuntek, H.; Epplen, J. T.; Schols, L.; Riess, O. Ala30Pro mutation in the gene encoding alpha-synuclein in Parkinson’s disease. Nat Genet 1998, 18 (2), 106–108. DOI: 10.1038/ng0298-106.

(8) Conway, K. A.; Lee, S. J.; Rochet, J. C.; Ding, T. T.; Williamson, R. E.; Lansbury, P. T., Jr. Acceleration of oligomerization, not fibrillization, is a shared property of both alpha-synuclein mutations linked to early-onset Parkinson’s disease: implications for pathogenesis and therapy. Proc Natl Acad Sci U S A 2000, 97 (2), 571–576. DOI: 10.1073/pnas.97.2.571.

(9) Jo, E.; McLaurin, J.; Yip, C. M.; St George-Hyslop, P.; Fraser, P. E. alpha-Synuclein membrane interactions and lipid specificity. J Biol Chem 2000, 275 (44), 34328–34334. DOI: 10.1074/jbc.M004345200.

(10) Giasson, B. I.; Murray, I. V.; Trojanowski, J. Q.; Lee, V. M. A hydrophobic stretch of 12 amino acid residues in the middle of alpha-synuclein is essential for filament assembly. J Biol Chem 2001, 276 (4), 2380–2386. DOI: 10.1074/jbc.M008919200.

(11) Sharon, R.; Bar-Joseph, I.; Frosch, M. P.; Walsh, D. M.; Hamilton, J. A.; Selkoe, D. J. The formation of highly soluble oligomers of alpha-synuclein is regulated by fatty acids and enhanced in Parkinson’s disease. Neuron 2003, 37 (4), 583–595. DOI: 10.1016/s0896-6273(03)00024-2.

(12) Suzuki, K.; Iseki, E.; Katsuse, O.; Yamaguchi, A.; Katsuyama, K.; Aoki, I.; Yamanaka, S.; Kosaka, K. Neuronal accumulation of alpha- and beta-synucleins in the brain of a GM2 gangliosidosis mouse model. Neuroreport 2003, 14 (4), 551–554. DOI: 10.1097/00001756-200303240-00004 From NLM Medline.

(13) Stockl, M.; Fischer, P.; Wanker, E.; Herrmann, A. Alphasynuclein selectively binds to anionic phospholipids embedded in liquid-disordered domains. J Mol Biol 2008, 375 (5), 1394–1404. DOI: 10.1016/j.jmb.2007.11.051 From NLM Medline.

(14) Vogiatzi, T.; Xilouri, M.; Vekrellis, K.; Stefanis, L. Wild type alpha-synuclein is degraded by chaperone-mediated autophagy and macroautophagy in neuronal cells. J. Biol. Chem. 2008, 283, 23542–23556.

(15) Kjaer, L.; Giehm, L.; Heimburg, T.; Otzen, D. The influence of vesicle size and composition on alpha-synuclein structure and stability. Biophys J 2009, 96 (7), 2857–2870. DOI: 10.1016/j.bpj.2008.12.3940 From NLM Medline.

(16) Parihar, M. S.; Parihar, A.; Fujita, M.; Hashimoto, M.; Ghafourifar, P. Alpha-synuclein overexpression and aggregation exacerbates impairment of mitochondrial functions by augmenting oxidative stress in human neuroblastoma cells. Int J Biochem Cell Biol 2009, 41 (10), 2015–2024. DOI: 10.1016/j.biocel.2009.05.008.

(17) van Rooijen, B. D.; Claessens, M. M.; Subramaniam, V. Lipid bilayer disruption by oligomeric alpha-synuclein depends on bilayer charge and accessibility of the hydrophobic core. Biochim Biophys Acta 2009, 1788 (6), 1271–1278. DOI: 10.1016/j.bbamem.2009.03.010 From NLM Medline.

(18) Auluck, P. K.; Caraveo, G.; Lindquist, S. alpha-Synuclein: membrane interactions and toxicity in Parkinson’s disease. Annu Rev Cell Dev Biol 2010, 26, 211–233. DOI: 10.1146/annurev.cellbio.042308.113313.

(19) Middleton, E. R.; Rhoades, E. Effects of curvature and composition on alpha-synuclein binding to lipid vesicles. Biophys J 2010, 99 (7), 2279–2288. DOI: 10.1016/j.bpj.2010.07.056 From NLM Medline.

(20) Ruiperez, V.; Darios, F.; Davletov, B. Alpha-synuclein, lipids and Parkinson’s disease. Prog Lipid Res 2010, 49 (4), 420–428. DOI: 10.1016/j.plipres.2010.05.004.

(21) van Rooijen, B. D.; Claessens, M. M.; Subramaniam, V. Membrane Permeabilization by Oligomeric alpha-Synuclein: In Search of the Mechanism. PLoS One 2010, 5 (12), e14292. DOI: 10.1371/journal.pone.0014292.

(22) Nasstrom, T.; Fagerqvist, T.; Barbu, M.; Karlsson, M.; Nikolajeff, F.; Kasrayan, A.; Ekberg, M.; Lannfelt, L.; Ingelsson, M.; Bergstrom, J. The lipid peroxidation products 4-oxo-2-nonenal and 4-hydroxy-2-nonenal promote the formation of alpha-synuclein oligomers with distinct biochemical, morphological, and functional properties. Free Radic Biol Med 2011, 50 (3), 428–437. DOI: 10.1016/j.freeradbiomed.2010.11.027.

(23) Shvadchak, V. V.; Falomir-Lockhart, L. J.; Yushchenko, D. A.; Jovin, T. M. Specificity and kinetics of alpha-synuclein binding to model membranes determined with fluorescent excited state intramolecular proton transfer (ESIPT) probe. J Biol Chem 2011, 286 (15), 13023–13032. DOI: 10.1074/jbc.M110.204776 From NLM Medline.

(24) Zhaliazka K.; Ali, A.; Kurouski, D. Phospholipids and Cholesterol Determine Molecular Mechanisms of Cytotoxicity of alpha-Synuclein Oligomers and Fibrils. ACS Chem Neurosci 2024, 15 (2), 371–381. DOI: 10.1021/acschemneuro.3c00671 From NLM Medline.

(25) Matveyenka, M.; Ali, A.; Mitchell, C. L.; Brown, H. C.; Kurouski, D. Cholesterol Accelerates Aggregation of alpha-Synuclein Simultaneously Increasing the Toxicity of Amyloid Fibrils. ACS Chem Neurosci 2024, 15 (21), 4075–4081. DOI: 10.1021/acschemneuro.4c00501 From NLM Medline.

(26) Matveyenka, M.; Ali, A.; Mitchell, C. L.; Sholukh, M.; Kurouski, D. Elucidation of cytotoxicity of alpha-Synuclein fibrils on immune cells. Biochim Biophys Acta Proteins Proteom 2024, 1873 (2), 141061. DOI: 10.1016/j.bbapap.2024.141061 From NLM Publisher.

(27) Asai, H.; Ikezu, S.; Tsunoda, S.; Medalla, M.; Luebke, J.; Haydar, T.; Wolozin, B.; Butovsky, O.; Kugler, S.; Ikezu, T. Depletion of microglia and inhibition of exosome synthesis halt tau propagation. Nat Neurosci 2015, 18 (11), 1584–1593. DOI: 10.1038/nn.4132.

(28) Yoshiyama, Y.; Higuchi, M.; Zhang, B.; Huang, S. M.; Iwata, N.; Saido, T. C.; Maeda, J.; Suhara, T.; Trojanowski, J. Q.; Lee, V. M. Synapse loss and microglial activation precede tangles in a P301S tauopathy mouse model. Neuron 2007, 53 (3), 337–351. DOI: 10.1016/j.neuron.2007.01.010 From NLM Medline.

(29) Maphis, N.; Xu, G.; Kokiko-Cochran, O. N.; Jiang, S.; Cardona, A.; Ransohoff, R. M.; Lamb, B. T.; Bhaskar, K. Reactive microglia drive tau pathology and contribute to the spreading of pathological tau in the brain. Brain 2015, 138 (Pt 6), 1738–1755. DOI: 10.1093/brain/awv081 From NLM Medline.

(30) Delamarre, L.; Pack, M.; Chang, H.; Mellman, I.; Trombetta, E. S. Differential lysosomal proteolysis in antigen-presenting cells determines antigen fate. Science 2005, 307 (5715), 1630–1634. DOI: 10.1126/science.1108003 From NLM Medline.

(31) Li, Q.; Barres, B. A. Microglia and macrophages in brain homeostasis and disease. Nat Rev Immunol 2018, 18 (4), 225–242. DOI: 10.1038/nri.2017.125 From NLM Medline.

(32) Liu, K. Dendritic Cells. Encyclopedia of Cell Biology 2016, 741–749.

(33) Michelucci, A.; Heurtaux, T.; Grandbarbe, L.; Morga, E.; Heuschling, P. Characterization of the microglial phenotype under specific pro-inflammatory and anti-inflammatory conditions: Effects of oligomeric and fibrillar amyloid-beta. J Neuroimmunol 2009, 210 (1-2), 3–12. DOI: 10.1016/j.jneuroim.2009.02.003 From NLM Medline.

(34) Mitchell, C. L.; Matveyenka, M.; Kurouski, D. Neuroprotective properties of transition metal dichalcogenide nanoflowers alleviate acute and chronic neurological conditions linked to mitochondrial dysfunction. J Biol Chem 2025, 301 (5), 108498. DOI: 10.1016/j.jbc.2025.108498 From NLM Medline.

(35) Jiang, Y.; Kang, Y.; Liu, J.; Yin, S.; Huang, Z.; Shao, L. Nanomaterials alleviating redox stress in neurological diseases: mechanisms and applications. J Nanobiotechnology 2022, 20 (1), 265. DOI: 10.1186/s12951-022-01434-5 From NLM Medline.

(36) Chen, T.; Zou, H.; Wu, X.; Liu, C.; Situ, B.; Zheng, L.; Yang, G. Nanozymatic Antioxidant System Based on MoS(2) Nanosheets. ACS Appl Mater Interfaces 2018, 10 (15), 12453–12462. DOI: 10.1021/acsami.8b01245 From NLM Medline.

(37) Han, Q.; Cai, S.; Yang, L.; Wang, X.; Qi, C.; Yang, R.; Wang, C. Molybdenum Disulfide Nanoparticles as Multifunctional Inhibitors against Alzheimer’s Disease. ACS Appl Mater Interfaces 2017, 9 (25), 21116–21123. DOI: 10.1021/acsami.7b03816 From NLM Medline.

(38) Ali, A.; Holman, A. P.; Rodriguez, A.; Matveyenka, M.; Kurouski, D. Tubulin-binding region alters tau-lipid interactions and changes toxicity of tau fibrils formed in the presence of phosphatidylserine lipids. Protein Sci 2024, 33 (7), e5078. DOI: 10.1002/pro.5078 From NLM Medline.

(39) Ali, A.; Holman, A. P.; Rodriguez, A.; Osborne, L.; Kurouski, D. Elucidating the mechanisms of alpha-Synuclein-lipid interactions using site-directed mutagenesis. Neurobiol Dis 2024, 198, 106553. DOI: 10.1016/j.nbd.2024.106553 From NLM Medline.

(40) Ali, A.; Matveyenka, M.; Pickett, D. N.; Rodriguez, A.; Kurouski, D. Tubulin-Binding Region Modulates Cholesterol-Triggered Aggregation of Tau Proteins. J Neurochem 2025, 169 (1), e16294. DOI: 10.1111/jnc.16294 From NLM Medline.

(41) Zhaliazka, K.; Kurouski, D. Elucidation of molecular mechanisms by which amyloid beta(1)(-)(42) fibrils exert cell toxicity. Biochim Biophys Acta Mol Cell Biol Lipids 2024, 1869 (6), 159510. DOI: 10.1016/j.bbalip.2024.159510 From NLM Medline.

