## Supplemental Figure 1 for "Transition Metal Dichalcogenide Nanoflowers Rescue Immune Cells from the Cytotoxic Effects of Amyloid Aggregates"

### Supporting Information

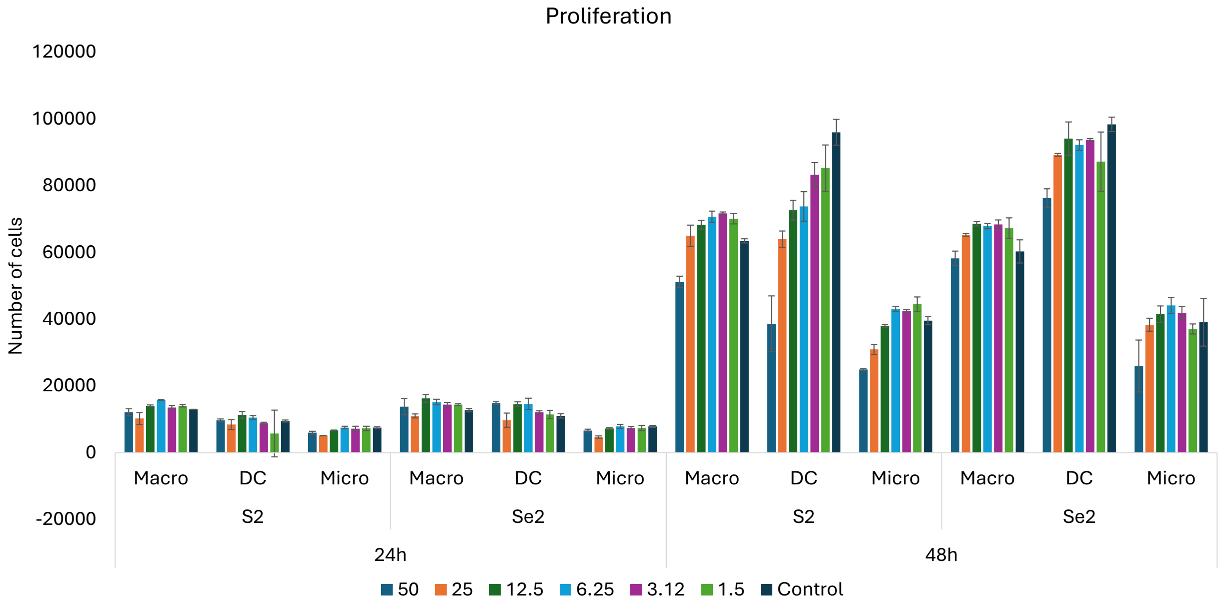


Figure S1. Proliferation of macrophages (macro), DC cells (DC) and microglia (micro) in the presence of 1.5-50 μM of $MoS_2$ (S2) and $MoSe_2$ (Se2).
